# Stem photosynthesis is coordinated with seasonal growth activity in two temperate tree species

**DOI:** 10.64898/2026.02.28.708753

**Authors:** Radek Jupa, Eva Harudová, Lenka Plavcová, Roman Plichta

## Abstract

Woody stems conduct both photosynthetic assimilation and respiration. The two processes work in concert, as stem photosynthesis helps refix CO_2_ released by stem respiration, thereby increasing carbon-use efficiency and generating a local pool of non-structural carbohydrates supporting cambial growth and stem hydraulic function. Despite its importance, little is known about seasonal variation in stem photosynthesis and the factors underlying its activity throughout the season. To fill this gap, we measured stem gas exchange together with growth activity, water status and photosynthetic pigment contents in two temperate species, *Acer platanoides* L. and *Prunus avium* L., over the season. In both species, gross photosynthetic rates (Pg) and dark respiration (Rd) changed significantly over the season in a similar pattern, indicating strong coordination between the two processes. Both Pg and Rd reached the highest values in May, during the period of rapid leaf expansion and secondary growth, and declined later in the growing season. At each measurement date, Rd exceeded Pg, resulting in a net CO_2_ efflux from the stems. The seasonal changes in Pg and Rd translated into seasonal variability in relative refixation of CO_2_, ranging from 3 to 59% and gradually decreasing towards the end of the season. Additionally, the Pg corresponded with the tissue hydration and increased significantly with increasing stem water potential. In contrast, total chlorophyll content showed less pronounced seasonal variation and thus explained substantially lower seasonal variability in Pg, except for the chlorophyll a/b ratio, which changed dynamically over the season and reached a minimum during the peak of the growing season. Overall, our results reveal that stem photosynthesis varies seasonally in accord with stem growth and water status, while the chlorophyll content has a lower impact on the seasonal changes. These findings are important for our understanding of the carbon relations of trees.

## Introduction

While photosynthesis is not the main function of woody stems, both bark and wood tissues are capable of light-dependent photo-assimilation of carbon dioxide. Photosynthetically active stems often occur in drought-adapted succulents and shrubs (Ávila-Lovera et al. 2017). In these species, the stems are often green and possess an epidermis with stomata. The photosynthetic characteristics of such stems resemble those of leaves because CO_2_ is taken up from the atmosphere and plant gas exchange can be readily regulated by stomata closing and opening. However, stem photosynthesis takes place also in trees, particularly in branches and young stems that are photosynthetically active (Pfanz et al. 2002, Rosell et al. 2015). This is possible due to the presence of photosynthetic pigments and related photosynthetic machinery in the parenchyma cells within bark, xylem, and eventually the pith in the case of the youngest branches (Schmitz et al. 2012, Natale et al. 2023a). In woody stems, CO_2_ for photosynthetic production is mostly recycled from stem respiration (Salomón et al. 2024). The carbon-recycling stem photosynthesis thus substantially reduces respiratory carbon loss. Across species, carbon recycling appears to vary considerably, from 7% to 100% of respired CO_2_ (Ávila et al. 2014).

Stem photosynthesis contributes a substantial portion of whole-plant carbon gain. For instance, stem photosynthesis accounted for 11% of wood production in branches of *Eucalyptus miniata* (Cernusak and Hutley 2011). The substantial contribution of stem photosynthesis to wood production was even higher (20-56%) in three evergreen woody angiosperms native to arid regions of California (Saveyn et al. 2010). The relative importance of stem photosynthesis could increase when leaf assimilation and phloem transport are diminished. This can occur during a period of drought-induced stomatal closure (Natale et al. 2023b) or due to leaf abscission, as was the case with drought-deciduous shrubs (Ávila-Lovera et al. 2017). In regions with temperature-dependent winter dormancy, stem photosynthesis is expected to be important during the spring season, when leaf area is not yet fully developed and light penetration into the canopy is relatively high (Urban et al. 2014). This can be particularly relevant in species that flower before the leaf flush since flowering is an energy-demanding process (Hoch et al. 2003).

More recently, stem photosynthesis has been linked to the maintenance of xylem hydraulic function (e.g., Trifilò et al. 2021, Natale et al. 2023b). By producing a local carbohydrate pool, stem photosynthesis provides readily available resources to support xylem-associated processes. Osmotically active sugars derived from stem photosynthesis can be released from the vessel-associated cells (VACs) and drive embolism reversal (Secchi and Zwieniecki 2011, Nardini et al. 2011, Schmitz et al. 2012). In addition, inhibition of stem photosynthesis by stem shading significantly increased xylem vulnerability to embolism (De Baerdemaeker et al. 2017, Natale et al. 2023b). The exact mechanism by which stem shading increases xylem vulnerability is not known. It is possible that the lack of sugars may lower turgor of VACs and thereby facilitate air-seeding at the VACs-vessel interface (Natale et al. 2023b) or reduce the synthesis of surfactants that stabilize nanobubbles against expansion (De Baerdemaeker et al. 2017, Schenk et al. 2017). Stem photosynthesis could also facilitate water uptake through bark and thereby contribute to the refilling of embolized vessels (Liu et al. 2019). Thus, stem photosynthesis may play an important role in tree recovery after drought stress. However, this mechanism is yet to be tested on a seasonal timescale.

There is sporadic evidence that stem photosynthesis changes seasonally. The seasonal variability could reflect the balance between changing demands for carbon assimilation (e.g., actual growth rate and tissue development), external environmental factors (e.g., light and water availability), and internal physiological factors (e.g., phenological stage and associated chlorophyll content) that influence photosynthetic production. For instance, across 14 succulent shrub species from several arid locations in California, stem photosynthesis was, on average, higher during the dry season than during the wet season (Ávila-Lovera et al. 2017). During the dry season, the shrubs were leafless and the light conditions within the canopy were more favorable for stem photosynthesis because of less shading and clearer skies. In nine temperate tree species, stem gross photosynthesis was lower in winter compared to summer and this trend was paralleled by a lower chlorophyll content and lower efficiency of photosystem II (Berveiller et al. 2007). However, measurements with a finer seasonal resolution are needed to better understand the importance of stem photosynthesis for the overall carbon budget and to elucidate the factors driving the photosynthetic rates along with the tissue development.

In this study, we observed seasonal variation in stem photosynthesis and dark respiration in relation to stem water status, photosynthetic pigment content, and radial growth progression in juvenile branches of two temperate tree species (*Acer platanoides* L. and *Prunus avium* L.). We hypothesized that stem photosynthesis and chlorophyll content would be highest during the peak of the cambial growth because carbohydrates are needed to support cell division and expansion. We also expected that stem photosynthetic rates would reflect stem water status, providing carbohydrates for maintaining xylem hydraulic function.

## Materials and Methods

### Site and plant material

The experiments were conducted on two tree species, *Acer platanoides* L. and *Prunus avium* L., growing at the Kamenný vrch site (Brno, Czech Republic; 49.1875633 N, 16.5509383 E; 370 m a.s.l.). The mean annual temperature at the site is 10.4 °C, and the mean annual precipitation is 531 mm. Broadleaved temperate trees and shrubs occur sparsely at the site, characterized by shallow cambisol with frequently exposed bedrock.

For each species, we randomly selected eight mature, healthy individuals growing within a 1.5-ha area. We primarily chose solitary trees fully exposed to sunlight. All selected individuals were more than 6 m tall, with average heights of 9 m for *A. platanoides* and 8 m for *P. avium*. Their size allowed multiple branch collections throughout the season without significantly reducing their canopy volume, which could influence both foliar and stem photosynthesis (Zheng et al. 2021).

### Climatic factors

Transmitted global irradiance, air temperature, and relative air humidity were continuously measured at 15-minute intervals using the Minikin RTHi datalogger (EMS Brno, Czech Republic). The datalogger was placed in a canopy margin of one of the trees 2 m above ground, corresponding to the position of the sampled branches. Vapor pressure deficit (VPD, kPa) was calculated according to Allen et al. (1998). For air temperature, daily mean values were calculated, while transmitted global irradiance and VPD were averaged for each day from measurements taken between 10 a.m. and 2 p.m. to reflect midday conditions. During the sampling dates, soil samples were taken at three different places within the locality at 10 cm depths and transferred to the laboratory in ziplocks, where soil water potential (ψ_S_, MPa) was determined with the dewpoint meter WP4C.

### Branch sampling and processing

In the sun-exposed southern canopy of each tree, we randomly selected seven branches longer than 1 m, located 2–3 m above ground. These branches were sampled across the prominent phenological phases from March to December 2024: 1) beginning of bud swelling (7^th^ March), 2) flowering and early leaf development (7^th^ April), 3) vegetative growth and fruit development initiation (16^th^ May), 4) maximum vegetative growth rate, continuous fruit development (20^th^ June), 5) vegetative growth reduction, finishing fruit development (24^th^ July), 6) vegetative growth cessation, leaf senescence initiation (20^th^ September), 7) end of leaf fall, dormancy initiation (5^th^ December; Figure 1). The phenological phases within each sampling date were assigned to the standardized BBCH scale according to Finn et al. (2007).

**Figure 1.**
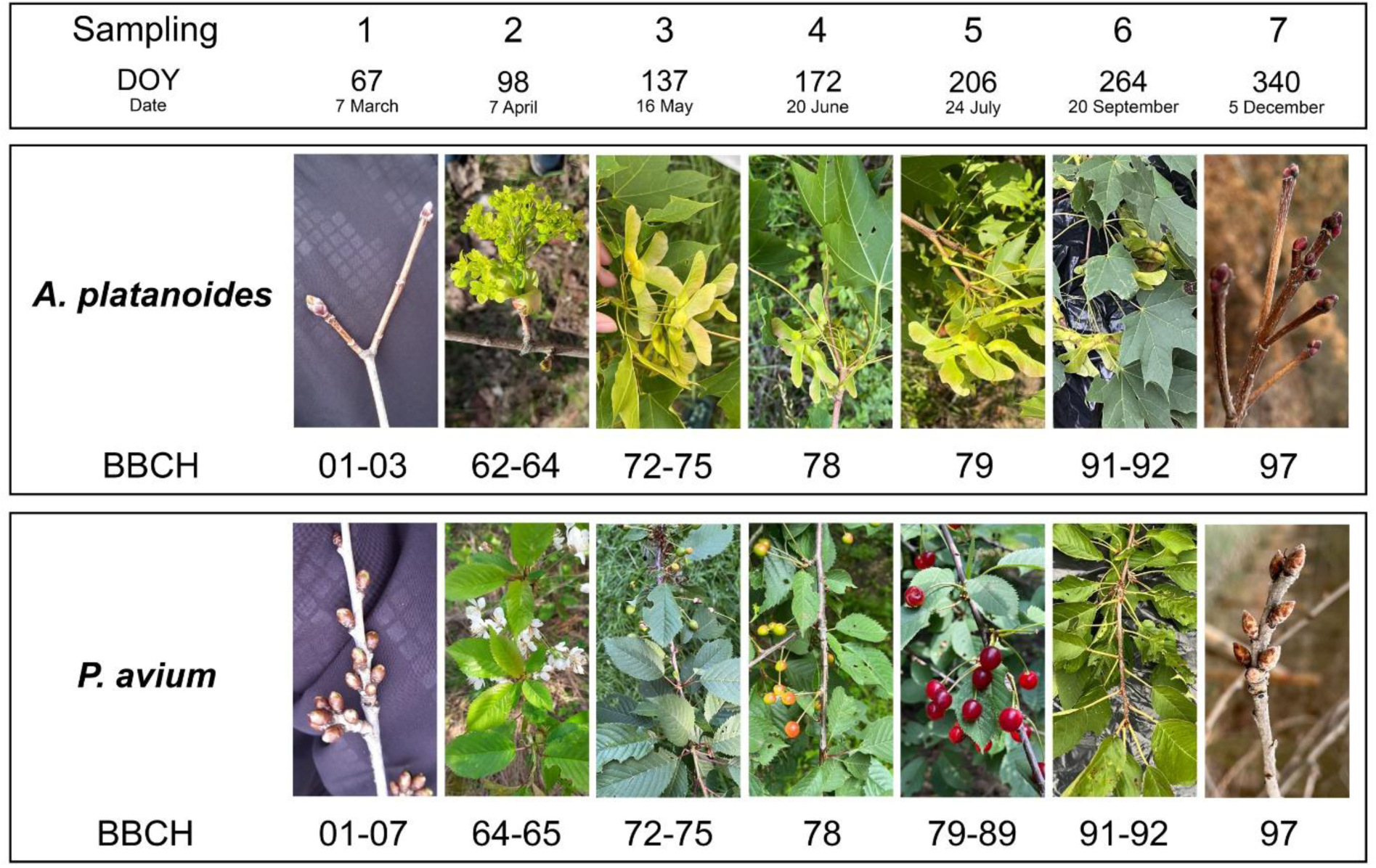
Typical phenological phases of *Acer platanoides* and *Prunus avium* recorded at individual sampling dates. The phenological phases were assigned to the standardized BBCH scale according to Finn et al. (2007).

At each sampling date, a single one-meter-long branch was collected from each individual tree during morning hours (7-8 a.m.). Initially, all leaves (if present) were cut off to minimize branch dehydration. To assess the effects of leaf area on photosynthetic activity and photoasimilation pigment contents, we collected all leaves associated with the branch segments used for gas exchange measurements, anatomical analyses and pigment contents (i.e., from the base of the current-year shoot to a distance of 45 cm basipetally from this point; Figure 2). The collected leaves were stored in a ziplock bag, and the total leaf area of each sample was subsequently measured using a portable scanner. The branch was then cut one meter from the apex, placed into a dark plastic bag, and immediately transported to the laboratory.

**Figure 2.**
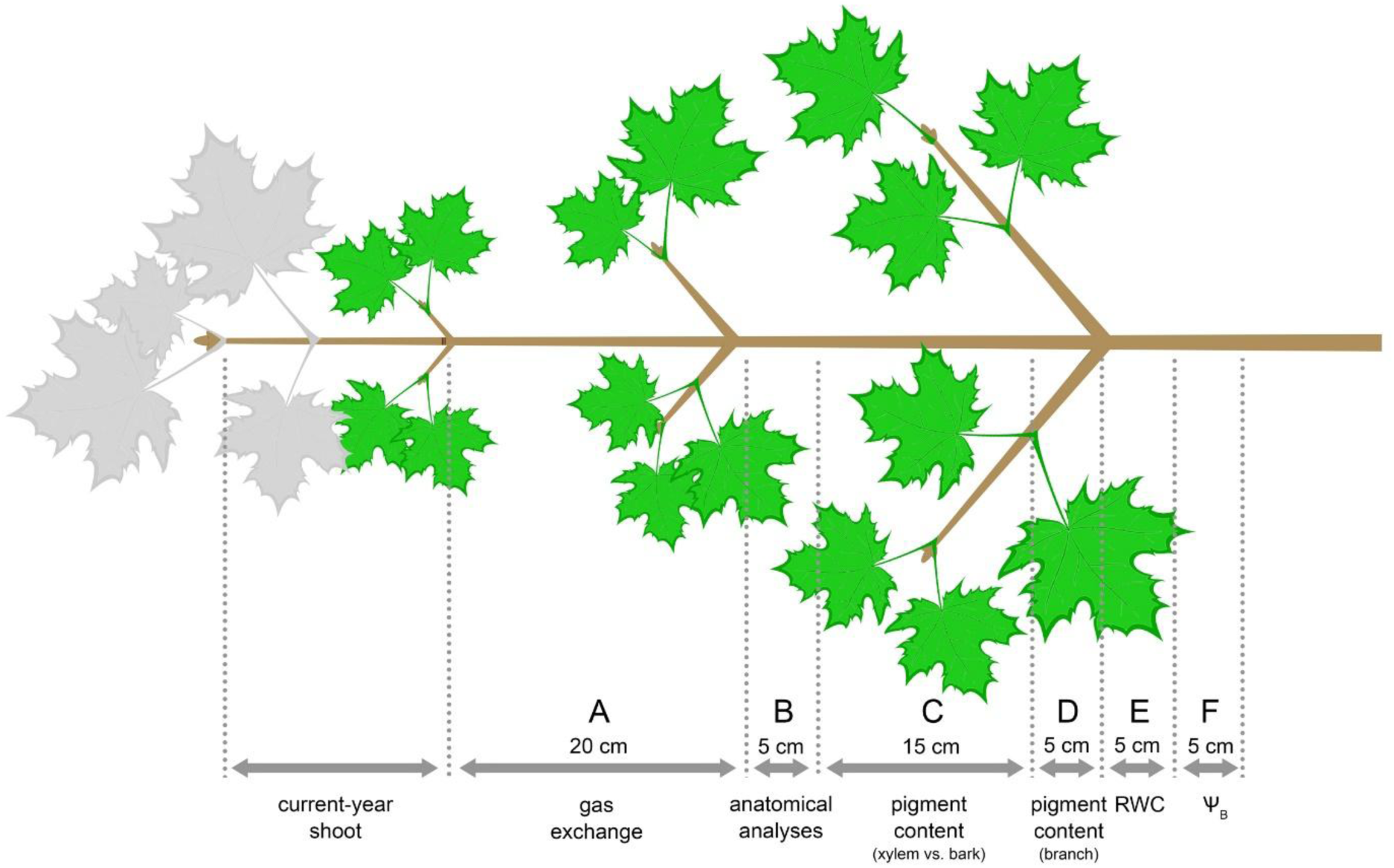
Scheme showing the processing of the sampled branch for analyses. Leaves used for leaf area determination are shown in green.

In the laboratory, the branches were cut into a series of individual segments that were used for analyses described below (Figure 2). First, the current-year shoot was removed and not used in any analyses. Segment A was excised between 0 cm and 20 cm from the current-year shoot base (SB) and used for gas exchange measurements (Figure 2). Segment B was excised between 20 cm and 25 cm from SB and used for the anatomical analyses. Segment C was excised between 25 cm and 40 cm from SB and used to determine photoasimilation pigment contents in xylem and bark. Segment D was excised between 40 cm and 45 cm from SB and used to determine photoasimilation pigment contents in the whole branch. Segment E was excised between 45 cm and 50 cm from SB and used to determine branch relative water content. Segment F was excised between 50 cm and 55 cm from SB and used to determine branch water potential. The age of all excised segments was 2-3 years. All measurements were done no later than six hours after branch sampling.

### Stem gas exchange

Immediately after excision, segment A was divided into two 10 cm-long segments (proximal and distal). Both cut ends were sealed with Parafilm to avoid excessive dehydration. The maximum net photosynthetic rate (Pn, µmol m^-2^ s^-1^) of both segments together was measured using the IRGA CO_2_ Analyzer Qubit model no. S151 (Qubit Systems, Canada) equipped with a custom-built chamber at the following conditions: reference CO_2_ concentration 450 ppm, irradiance 2000 µmol m^-2^ s^-1^, flow rate 15 l hod^-1^and air temperature 25°C. After reaching steady-state values, the lights were switched off, and the chamber containing both segments was darkened to measure dark respiration (Rd, µmol m^-2^ s^-1^). The duration of gas exchange measurements typically did not exceed 15 min. Subsequently, dimensions of both segments were precisely determined with a caliper, and the surface area of each segment (*A*, m^2^) was determined as the lateral surface area of the truncated cone. Both Pn and Rd were calculated as Pn(Rd)=(Δc × f × p)/(R × T × A), where Δc is the difference in CO_2_ concentration between reference and the steady state value, *f* is the flow rate, *p* is the atmospheric pressure (Pa), *R* is the universal gas constant (J K^-1^ mol ^-1^), and *T* is the absolute temperature of the air in the chamber (K). The maximum gross photosynthetic rate (Pg, µmol m^-2^ s^-1^) was then determined as Pg=Pn+Rd. Relative CO_2_ refixation (Refix, %) was calculated according to Damesin (2003) as Refix=[(Rd – Pn)/Rd]*100.

### Anatomical analyses

Segment B was shortened and divided into quarters longitudinally with a sharp razor blade. The segment quarters were stored in FAA fixative until further processing. For each branch, permanent slides were prepared from one randomly selected quarter as described by Ruzin (1999). Specifically, the samples were continuously dehydrated in an increasing butanol series and embedded in high melt paraffin (Paraplast, Leica, Germany). From the samples, 15 µm-thick cross-sections were prepared using a Reichert sledge microtome, dewaxed in xylene, and stained with 1% (w/v) aqueous Safranin solution, which stains lignified cell walls red. The sections were mounted on slides with Eukitt (Sigma-Aldrich), observed in bright field using an Olympus BX-50 optical microscope (Olympus, Japan) at 200× magnification, and photographed with the attached Sony A35 camera (Sony, Japan). The photographs were merged into a single image using Adobe Photoshop CS6 (Adobe, USA), and the thicknesses of the xylem and bark layers were determined in Fiji software (ImageJ 2.1.0/1.53c; Schindelin et al. 2012). We analyzed the thicknesses of xylem (i.e., distance between pith margin and mid of the cambial zone), cambial zone (i.e., all cambium derivates having not fully developed cell walls), secondary phloem (i.e., distance between the mid of the cambial zone and the margin of the last ray cell), cortex (i.e., distance between secondary phloem and phellogen), and phellem (i.e., distance between phellogen and last phellem cell). The total bark thickness included secondary phloem, cortex, and periderm thicknesses. Parenchymatic phelloderm was not distinguished from the cortex. The thicknesses were measured within three regions of the branch quarter and averaged for each sample.

### Pigment contents

For analyses of photoassimilation pigment content, we followed the methodology described by Bloemen et al. (2016). Bark was detached from the xylem in segment C. Subsequently, the separated bark and xylem from segment C and the whole branch segment D were cut into small pieces with a sharp razor blade. The cut pieces were immediately frozen in liquid nitrogen and stored in a freezer until further processing. Frozen samples were lyophilized for 48 h and homogenized using a rotary mill (Retch ZM 100, Retch, Germany). The pigments were extracted from bark and whole branch samples by adding 7.5 ml of 80% aceton and 10 mg of MgCO_3_ to 150 mg of homogenized dry mass, while 350 mg of dry mass was used for the extraction of pigments from xylem samples due to substantially lower pigment contents. After 24 h of extraction on an orbital shaker in the dark, the samples were centrifuged (500G, 10 min), and 2.8 ml of supernatant was transferred to a glass cuvette. The absorptions were measured spectrophotometrically at 470 nm, 646.6 nm, and 663.6 nm using the Specor 205 spectrophotometer (Analytik Jena, Germany). Concentrations of chlorophyll a, chlorophyll b, and carotenoids were calculated according to Porra et al. (1989) and Wellburn (1994). The concentrations were converted to contents of chlorophyll a (Chl a, mg g^-1^), chlorophyll b (Chl b, mg g^-1^), and carotenoids (Car, mg g^-1^) normalized per unit of dry weight.

### Relative water content and water potential

Segment E was weighed immediately after its excision on an analytical balance (Radwag AS 220/X, Radwag, Poland) to determine its native weight (m_n_). Subsequently, the segment was saturated by water under vacuum overnight, and its saturated weight was determined (m_s_). The segment was then dried in a laboratory oven at 70°C for 72 h, and its dry weight was measured as described before (m_d_). The relative water content (RWC, %) was calculated as RWC=(m_n_-m_d_)/(m_s_-m_d_).

The whole segment F was cut into smaller pieces to determine branch water potential (ψ_B_, MPa) using a previously calibrated dewpoint meter WP4C (METER Group, USA).

### Statistical analyses

To evaluate the seasonal time-course of the measured variables, linear mixed-effect models were fitted with species and date as fixed effects and tree individual as a random effect, recognizing that measurements throughout the season were made repeatedly on the same individual trees. The fitting was done using lmer function from the lme4 package (Bates et al. 2015). Model assumptions, including the normality of residuals and homogeneity of variance, were verified through diagnostic plots. Following the fitting, type III ANOVA tables were generated to assess the significance of the fixed factors. *P*-values were calculated using Satterthwaite’s approximation for denominator degrees of freedom via the lmerTest package (Kuznetsova et al. 2017). Post hoc comparisons among sampling times within each species were conducted using estimated marginal means (EMMs) with the emmeans package (Lenth 2025). Compact letter displays were generated to indicate statistically significant differences between groups based on pairwise comparisons with Tukey-adjusted *p*-values. Linear mixed-effect models were also fitted to evaluate relationships between branch gas exchange variables and potential explanatory variables. As previously, the significance of the effects was evaluated using the type III ANOVA with the Satterthwaite’s approximation for denominator degrees of freedom. The results are presented in the form mean ± SD (text) and mean ± SE (graphs) unless stated otherwise. All analyses were conducted in R (version 4.5.0, R Core Team 2024).

## Results

### Microclimatic conditions

Both daily air temperature and midday VPD within the canopy followed a seasonal pattern typical of Central European climates (Figure 3a–b). From March onward, both variables showed an increasing tendency, reaching their highest daily values of approximately 26.5 °C and 3.5 kPa at the beginning of September. After September, air daily temperature and midday VPD declined rapidly, reaching seasonal minima of around 3 °C and 0 kPa in early December.

**Figure 3.**
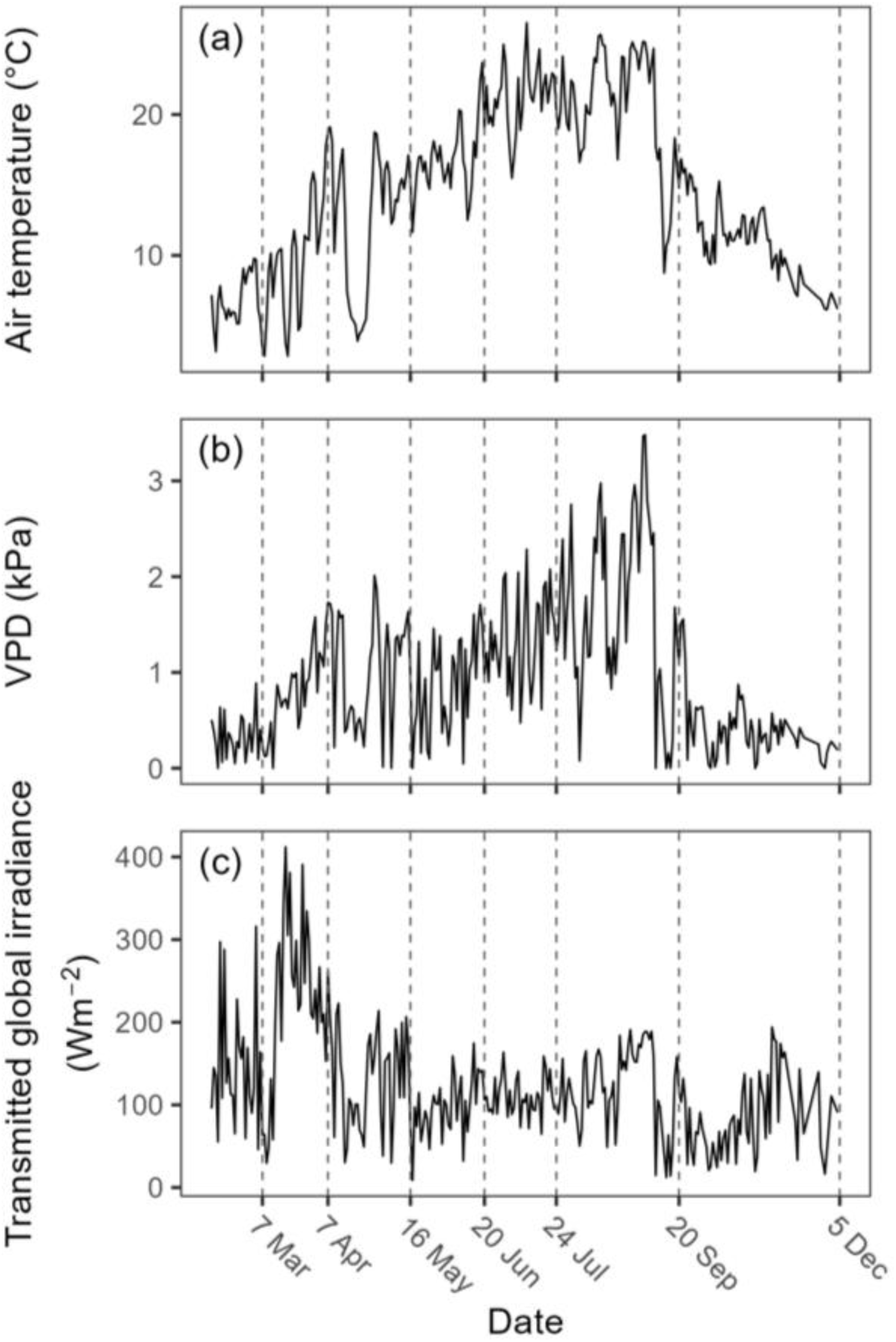
The courses of daily air temperature (a), midday vapor pressure deficit (b) and midday transmitted global irradiance (c) measured in a canopy of a selected tree during the year. The dashed vertical lines represent the sampling dates.

Midday transmitted global irradiance within the canopy ranged between 100 and 200 W m⁻² and remained relatively stable throughout the growing season, except during the pre-leaf-flush period in spring, when values were generally higher (Figure 3c).

Soil water potential in the upper soil layer remained relatively high across sampling dates and dropped below 0 MPa only in May (−0.12 MPa), July (−0.23 MPa), and December (−0.06 MPa; Figure S1).

### Secondary growth

The progression of branch secondary growth throughout the growing season was obvious from anatomical observations. The thickness of the cambial zone was the highest in May and June, corresponding to the phase of most active secondary growth (Figure 4a). As a result of progressive secondary growth, xylem thickness increased gradually till mid-summer in both species (Figure 4b), while the seasonal variation in bark thickness was less pronounced (Figure 4c). Within the bark tissues, the thickness of the phloem, cortex, and phellem showed more variable seasonal patterns (Figure 4d-f). In *A. platanoides*, phloem thickness increased throughout the growing season and decreased relatively sharply in December (Figure 4d), while cortex and phellem thickness showed a weak seasonal variation (Figure 4e-f). In *P. avium*, phloem thickness increased slightly during the growing season but did not decline in winter (Figure 4d). The cortex thickness showed only a weak seasonal variation (Figure 4e), while, in contrast to *A. platanoides*, phellem thickness increased sharply from May to June (Figure 4f).

**Figure 4:**
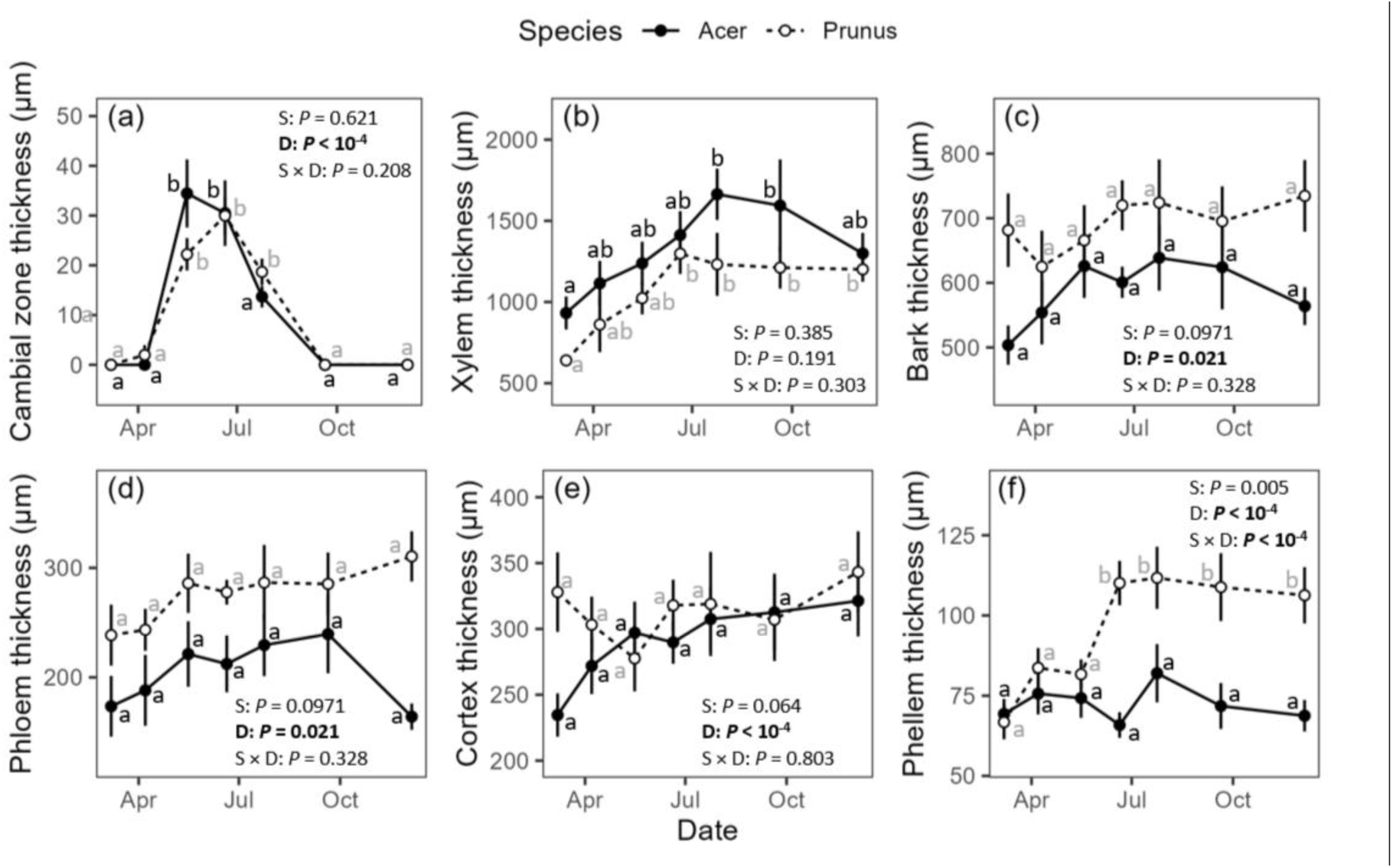
Seasonal variation in thicknesses of cambial zone (a), xylem (b), bark (c), phloem (d), cortex (e) and phellem (f) in *Acer platanoides* (closed circles) and *Prunus avium* (open circles). A linear mixed-effects model accounting for repeated measurements on individual trees was used to evaluate the effects of species (S), sampling date (D), and their interaction (S × D). Statistically significant fixed effects are highlighted in bold. Different letters indicate significant differences within individual species among the sampling dates. Data are in the form means ± SE (n=8).

### Branch water status

Branch relative water content (RWC) and branch water potential (Ψ_B_) showed a relatively weak seasonal pattern in both species (Figure 5a-b). However, there was a tendency to reach higher RWC and Ψ_B_ during the period of active growth between April and June (Figure 5a-b).

**Figure 5.**
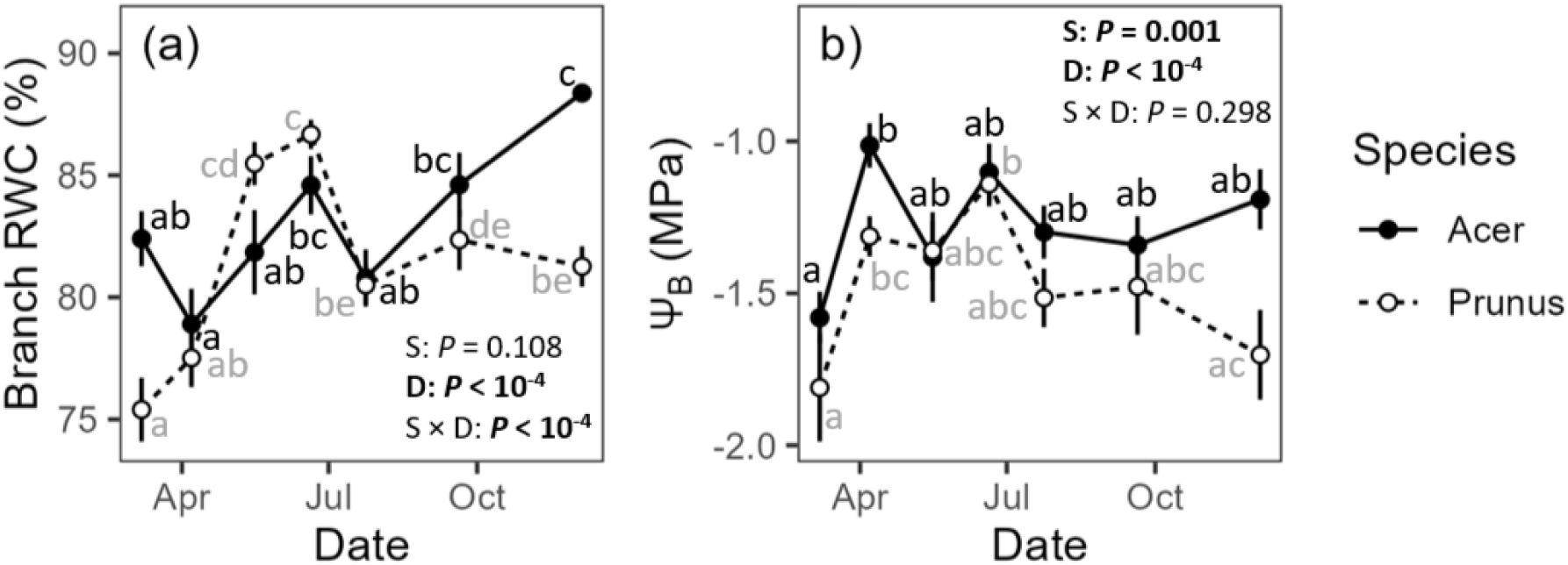
Seasonal variation in branch relative water content (a) and branch water potential (b) in *Acer platanoides* (closed circles) and *Prunus avium* (open circles). A linear mixed-effects model accounting for repeated measurements on individual trees was used to evaluate the effects of species (S), sampling date (D), and their interaction (S × D). Statistically significant fixed effects are highlighted in bold. Different letters indicate significant differences within individual species among the sampling dates. Data are in the form means ± SE (n=8).

### Pigment content

Total chlorophyll content in branches was generally higher in *P. avium* than in *A. platanoides* and showed a relatively weak seasonal variation with a slightly higher content between April and September (Figure 6a). In contrast to the relatively small seasonal change in total chlorophyll content in branches, there was a relatively stronger seasonal decline in chlorophyll a/b ratio in branches, with the minimal values in May, June, and July (Figure 6b). The seasonal patterns in the total chlorophyll contents and chlorophyll a/b ratio were mainly driven by the pigment contents in the bark and less so by those of the xylem (compare Figures 6c-d and e-f). The decline in the chlorophyll a/b ratio was mainly driven by the higher chlorophyll b content (compare Figures S2a, d, g and S2b, e, h). On average, bark contained 13 times and 6 times greater chlorophyll contents than xylem in *A. platanoides* and *P. avium*, respectively. Similar to chlorophyll contents, carotenoid contents in branches were more abundant in *P. avium* than *A. platanoides* (Figure S2c, f, i). The seasonal variation in carotenoids was weak in xylem, while increased carotenoid contents were measured in the branches and bark during the peak of the growing season.

**Figure 6:**
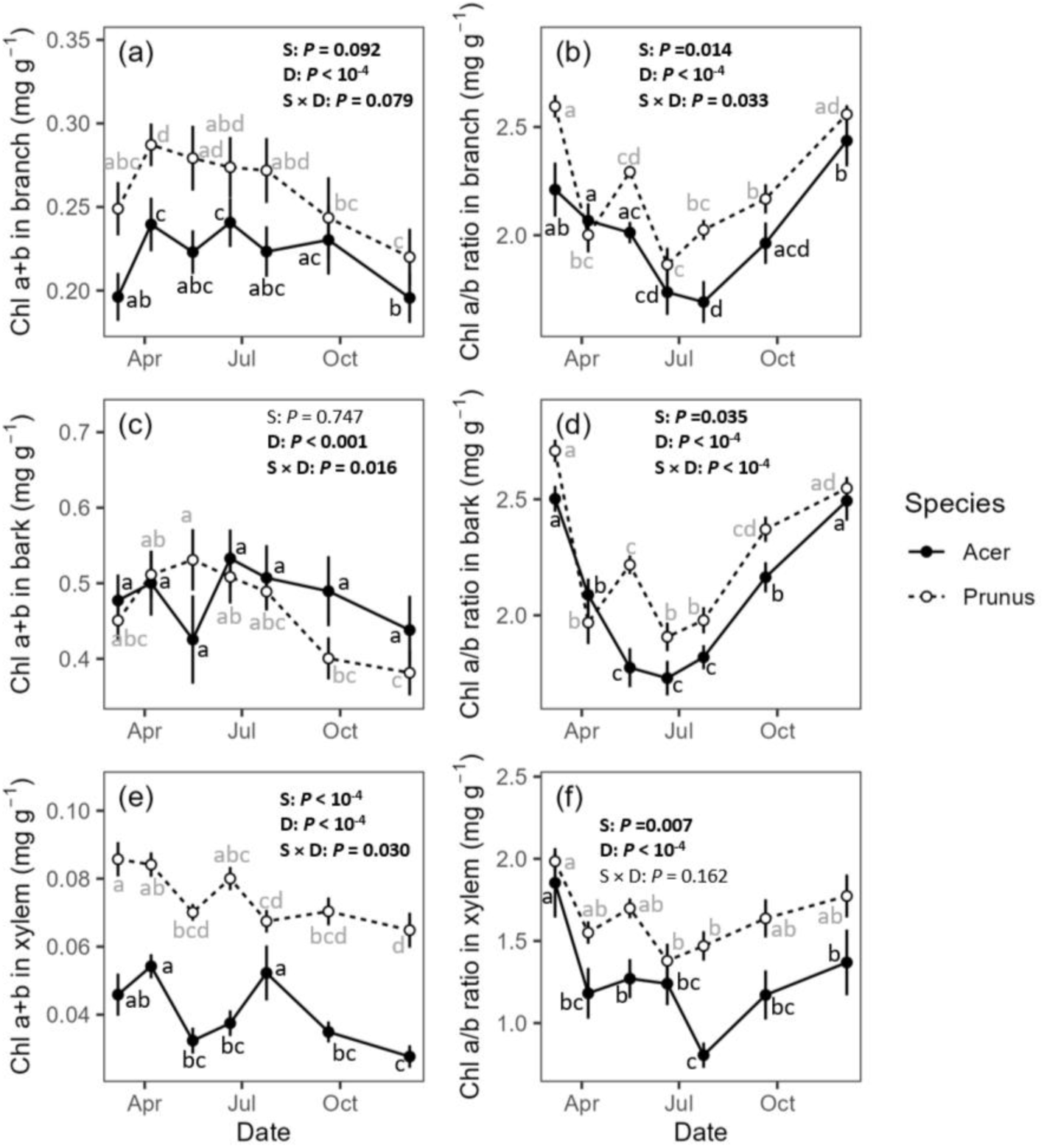
Seasonal variation in chlorophyll a + b contents in the whole branch (a), chlorophyll a/b ratio in the whole branch (b), chlorophyll a + b contents in bark (c), chlorophyll a/b ratio in bark (d), chlorophyll a + b contents in xylem (e), and chlorophyll a/b ratio in xylem (f) of *Acer platanoides* (closed circles) and *Prunus avium* (open circles). A linear mixed-effects model accounting for repeated measurements on individual trees was used to evaluate the effects of species (S), sampling date (D), and their interaction (S × D). Statistically significant fixed effects are highlighted in bold. Different letters indicate significant differences within individual species among the sampling dates. Data are in the form means ± SE (n=8).

### Branch gas exchange measurements

We observed significant changes in photosynthetic and dark respiration rates over the season (Figure 7). Net photosynthetic rates reached the lowest (i.e., most negative) values in May and the highest (i.e., least negative) values were measured in March or December (Figure 7a). Net photosynthetic rates were negative in both species during all measurement dates, indicating that the dark respiration rates (Figure 7b) were greater than gross photosynthetic rates (Figure 7c) throughout the season in both species. The dark respiration and gross photosynthetic rates were the highest in May and June (corresponding to the period of active growth) and decreased towards the end of the growing season. The maximum dark respiration rates were 2.46 ± 0.83 µmol m^-2^ s^-1^ and 3.13 ± 1.27 µmol m^-2^ s^-1^ in *A. platanoides* and *P. avium*, respectively. The gross photosynthetic rates showed greater interspecific differences and reached maximum values of 0.40 ± 0.22 µmol m^-2^ s^-1^ and 1.26 ± 0.68 µmol m^-2^ s^-1^ in *A. platanoides* and *P. avium*, respectively. The drop in both dark respiration and photosynthetic rates was steepest from June to July in *P. avium*, while the seasonal variation was lowest for gross photosynthetic rates in *A. platanoides*. Also, we observed seasonal variability in the relative CO_2_ refixation (Figure 7d). Relative refixation was highest in early spring before the onset of growth and decreased gradually towards the end of the growing season (Figure 7d). Similar to the gross photosynthetic rates, relative refixation was significantly lower in *A. platanoides* compared to *P. avium* over the whole season and varied between 3 and 36% in *A. pseudoplatanus* and 25 and 59% in *P. avium*.

**Figure 7:**
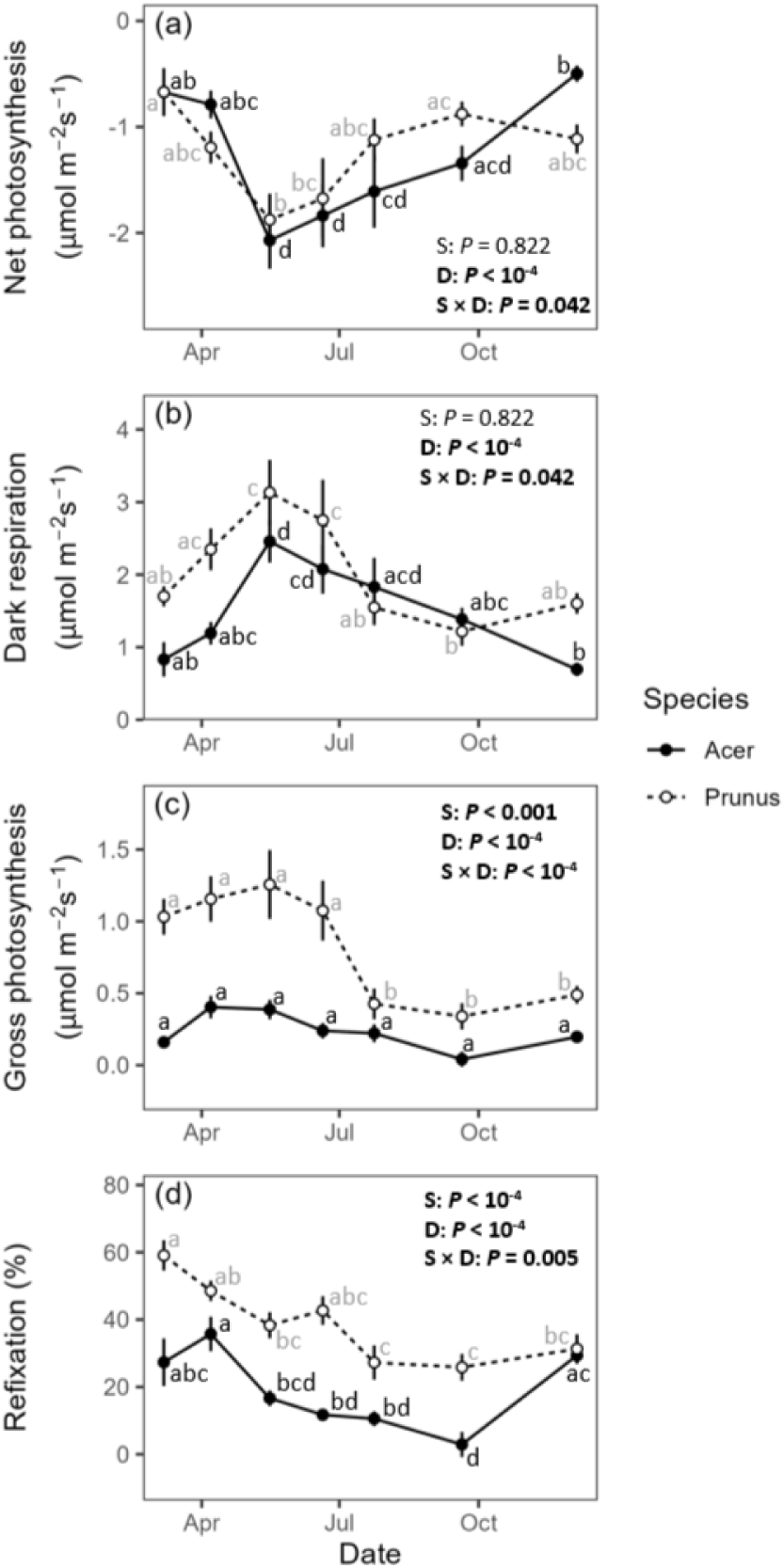
Seasonal variation of net photosynthetic rates (a), dark respiration rates (b), gross photosynthetic rates (c) and relative CO_2_ refixation (d) in branches of *Acer platanoides* (closed circles) and *Prunus avium* (open circles). Linear mixed effect model accounting for the repeated measurements on individual trees was used to evaluate the effect of species (S) and sampling date (D) and their interaction (S × D). Statistically significant fixed effects are highlighted in bold. Different letters indicate significant differences within individual species among the sampling dates. Data are in the form means ± SE (n=8).

### Relationships between gas exchange, physiological and anatomical traits

Gross photosynthetic rates were closely related to the dark respiration in both species (Figure 8), indicating a close coordination of both metabolic processes within the season. Moreover, gross photosynthetic rates were related to branch water potential, with greater values of gross photosynthetic rates being measured in branches with a less negative water potential (Figure 9a). Gross photosynthetic rates were not significantly related to total chlorophyll content of the branch (Figure 9b). Instead, gross photosynthetic rates were positively related to branch leaf area (Figure 9c). The latter relationship was steeper for *P. avium* than for *A. platanoides*.

**Figure 8:**
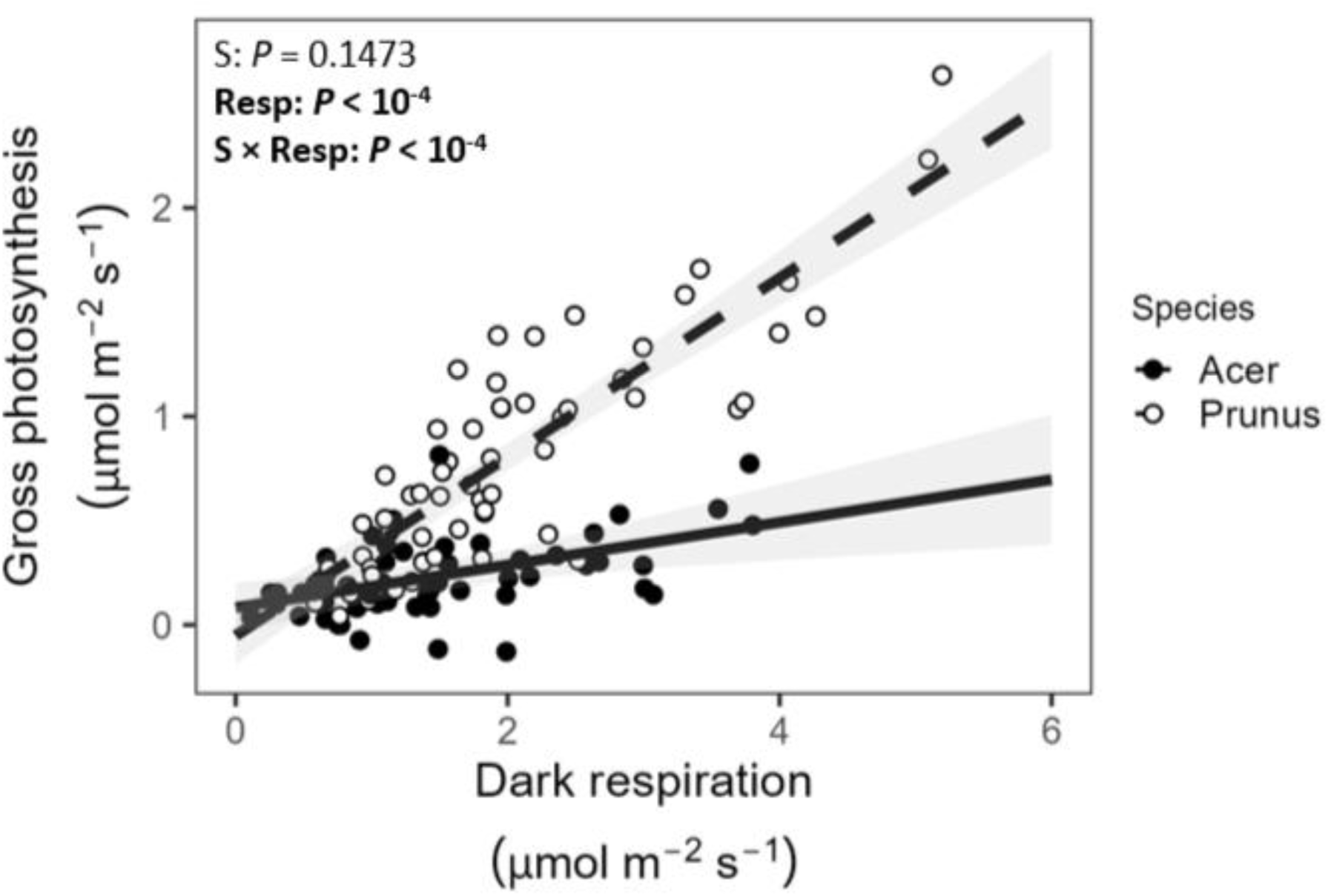
Relationship between dark respiration and gross photosynthetic rates in branches of *Acer platanoides* (closed circles, solid line) and *Prunus avium* (open circles, dashed line). A linear mixed-effects model accounting for repeated measurements on individual trees was used to evaluate the effects of species (S), respiration (Resp), and their interaction (S × Resp). Statistically significant fixed effects are highlighted in bold.

**Figure 9:**
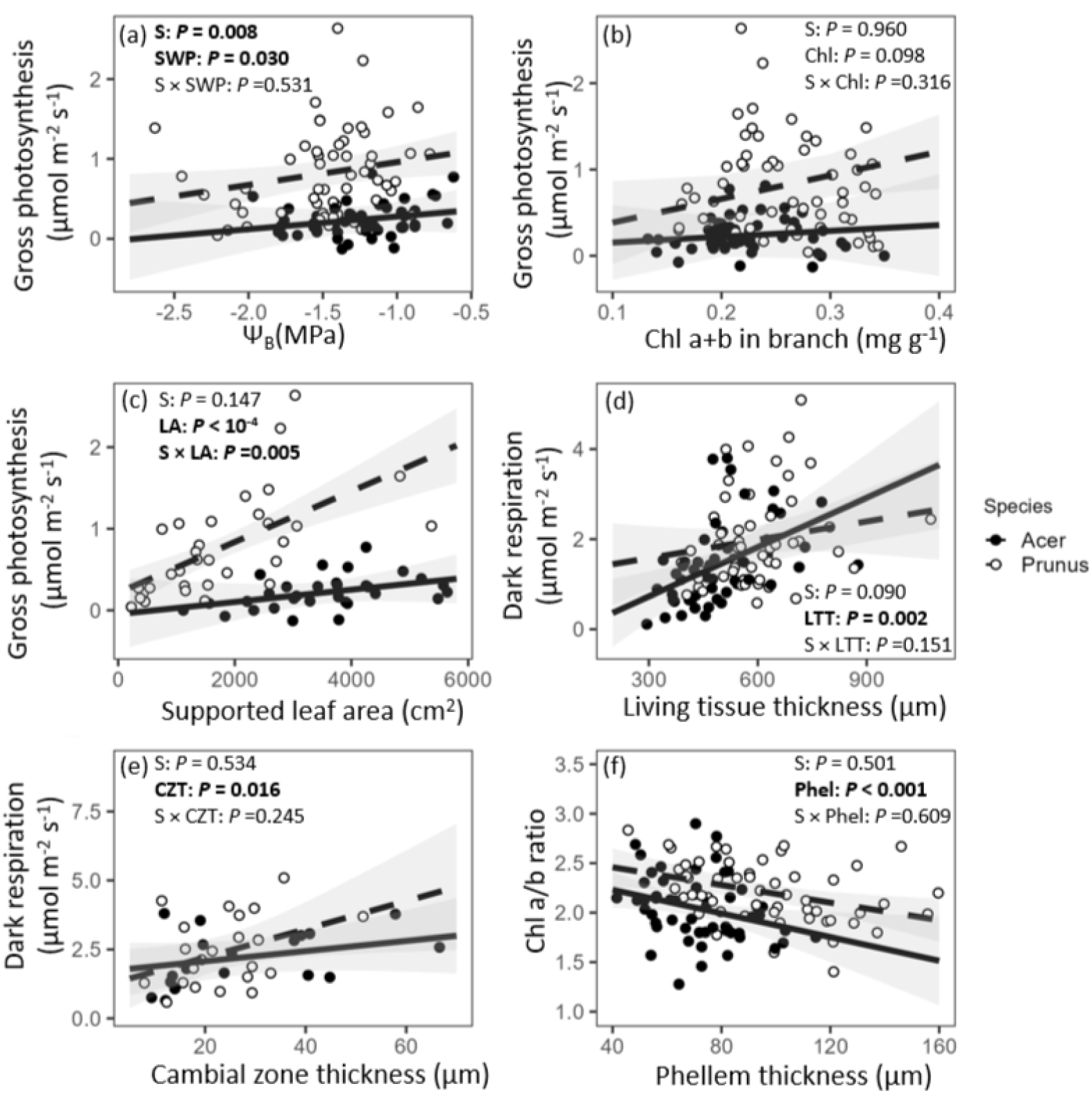
Relationships between gross photosynthetic rates and branch water potential (a), gross photosynthetic rates and chlorophyll a+b content in the whole branch (b) gross photosynthetic rates and leaf area (c), dark respiration rates and living tissue thickness (d), dark respiration rates and cambial zone thickness (e) and chlorophyll a/b ratio and phellem thickness (f) in branches of *Acer platanoides* (closed circles, solid line) and *Prunus avium* (open circles, dashed line). A linear mixed-effects model accounting for repeated measurements on individual trees was used to evaluate the effects of species (S), variable of interest and their interaction (S × variable abbreviation). Statistically significant fixed effects are highlighted in bold.

Dark respiration rates increased with increasing phloem, cortex and cambial zone thickness (i.e., living tissue thickness), representing the majority of actively growing and metabolically active cells in the studied branches (Figure 9d) as well as cambial zone thickness (Figure 9e). Interestingly, the ratio between chlorophyll a and b contents in the branch decreased with increasing phellem thickness in both species (Figure 9f).

## Discussion

Our study provides a detailed assessment of seasonal variation in stem respiration and photosynthesis in two temperate tree species, *Acer platanoides* and *Prunus avium*. As we found, dark respiration rates peaked during the transition from late spring to early summer and scaled positively with cambial zone thickness, reflecting the high energy demand associated with the division of cambial initials and the maturation of their derivatives. Respiration rates may also reflect phellogen activity, whose seasonal activity appears to parallel that of the vascular cambium (Brzostowska et al. 2025). The phellogen seasonal dynamics of the studied species was apparent from the increase in phellem thickness, which was observed between May and July, particularly in *P. avium*. However, no clear relationship was found between phellem thickness and stem respiration rates, which may reflect the relatively small proportion of cells produced by phellogen and, therefore, the lower energetic demand compared with the activity of vascular cambium. Respiration rates were also correlated with inner bark thickness, which represents the bulk of living cells in stems. In addition, both species contain living ray and axial parenchyma cells in the xylem and the xylem of *A. platanoides* also includes living fibers (Plavcová et al. 2016). All these cells carry out maintenance respiration and some of them, for instance, the companion cells in the phloem or vessel-associated parenchyma cells, are particularly metabolically active, as evidenced by their high abundance of mitochondria and strong activity of acid phosphatase, and therefore contribute substantially to the whole-stem respiration (DeWitt and Sussman 1995, Alves et al. 2001).

Due to prevailing dark respiration over the photosynthetic activity, both studied species exhibited negative net photosynthetic rates throughout the study period, indicating CO_2_ efflux from illuminated stems throughout the year. Similar to our results, negative net photosynthetic rates have also been reported in branches of several temperate tree species during summer and winter (Berveiller et al. 2007). These, as well as our results, show that the stem photosynthesis of temperate species may not meet the carbohydrate demands for respiration during the year. The carbohydrate demands of stems must be supplied by products of foliar photosynthesis via phloem flow or by the utilization of reserve non-structural carbohydrates. Nevertheless, the net CO_2_ efflux was partly diminished under light by stem photosynthesis that reassimilated on average 19 % and 39 % of CO_2_ released by the respiration in *A. platanoides* and *P. avium*, respectively. The CO_2_ refixation rate was highest in early spring and at the end of the growing season, supporting our hypothesis that stem photosynthesis is favored under leafless conditions, when stem irradiance is likely to be highest (Urban et al. 2014).

Accordingly, gross photosynthesis reached the highest values in both studied species during spring and early summer, suggesting its functional significance during the early phenological phases of seasonal development. The resumption of growth in the spring is highly dependent on the reserves of non-structural carbohydrates (Tixier et al. 2019). Depletion of carbohydrate reserves over winter may represent a significant limitation for the early phenological phases of development, such as bud and leaf development. The observed upregulation of stem photosynthesis early in the spring may help mitigate the loss of carbohydrate reserves and directly promote bud and leaf development. The significance of stem photosynthesis for bud development was documented by Damesin (2003), who observed a peak in gross photosynthetic rates during the bud burst in branches of *Fagus sylvatica*. In addition, stem photosynthesis was shown to support buds by supplying sugars, as demonstrated using carbon isotope discrimination in three defoliated tree species (Saveyn et al. 2010). Carbohydrates produced by stem photosynthesis in early spring may also be important for supporting precocious flowering in *P. avium*, followed by rapid fruit development. In contrast, *A. platanoides* has green flowers and fruits with a higher surface-to-volume ratio that may be more photosynthetically efficient, enabling them to contribute assimilates for their own growth and development, and thereby reducing the need for stem-derived carbohydrates early in the season (Hoch 2005). Later, during the onset of secondary growth extending into May in both species, elevated gross photosynthetic rates may support the production of new biomass, particularly wood, which may be important for the recovery of xylem transport capacity after winter dormancy in some species (Cernusak and Hutley 2011). In addition, it can be anticipated that stem photosynthesis could primarily contribute to phellogen activity and periderm formation because photosynthetically active tissue is located closer to the phellogen in the stem periphery compared to phloem, although direct evidence for this mechanism is currently lacking in the literature.

In both species, dark respiration and gross photosynthetic rates were positively correlated, indicating close synchrony between stem carbon source and sink processes throughout the season. The close relationship between dark respiration and gross photosynthetic rates was also observed in young individuals of five temperate angiosperm tree species, including *P. avium*, during the growing season (Wittmann and Pfanz 2008) and in branches of *Populus tremula* measured during a single summer month (Aschan et al. 2001). The linear relationship suggests that stem photosynthetic capacity increases when CO_2_ released by respiration is abundant and, conversely, that photosynthetic activity may decline as CO_2_ concentration within the stem decreases. Although the exact physiological mechanism is not clear, the close seasonal coordination between stem respiration and photosynthetic activity may be a result of an interlinked regulatory loop that helps maintain the balance in concentrations of CO_2_ and O_2_ within the stem.

Stem internal CO_2_ concentrations are typically significantly higher (up to 26%; Pfanz and Aschan 2001) while O_2_ concentrations are generally lower compared to the atmosphere (often below 10%; Spicer and Holbrook 2005). High respiration rates during the peak of the season may significantly increase CO_2_ and decrease O_2_ concentration, potentially leading to acidification of the cell cytoplasm and hypoxia, and resulting in reduced enzymatic activity and reduced metabolism (Wittmann and Pfanz 2014). The increased photosynthetic activity may thus remove excess CO_2_ and support cells with O_2,_ helping to avoid hypoxia. These effects may be particularly relevant to the highly metabolically active stem regions, such as the cambial zone and phloem. Besides, linking respiration and photosynthetic rates in the stem may help stabilize the carbohydrate content available. Increased gross photosynthetic rates may provide carbohydrates to support respiratory processes, while the CO_2_ released during respiration can stimulate photosynthesis.

Surprisingly, stem photosynthesis was not associated with changes in stem chlorophyll content throughout the season. This was likely due to the weak seasonal pattern of chlorophyll content, which decreased slightly toward the end of the season and reached a minimum during dormancy. While the low chlorophyll content observed during dormancy is more likely temperature-driven (Ivanov et al. 2006), a slight decrease during the season could partly reflect a gradual allocation of biomass to branch tissues (Cuny et al. 2015), affecting dry-matter-specific chlorophyll content. Decreasing dry-matter chlorophyll content during the growing season has also been reported for branch tissues of *F. sylvatica*, whereas area-based content showed almost no change at the same period (Damesin 2003). The higher photosynthetic rates in *P. avium* could be partially attributed to its higher branch chlorophyll content. In contrast to branch chlorophyll content, leaf chlorophyll typically shows more pronounced seasonal dynamics, with higher contents in summer and lower contents in spring and autumn (Demarez 1999, Ivanov et al. 2006, Croft et al. 2017). The weaker seasonal pattern in total branch chlorophyll contents is consistent with the perennial nature of stem tissue, in contrast to the annual cycle of leaf flushing and abscission. Regardless of the chlorophyll content, stem gross photosynthetic rates scaled positively with leaf area supported by the stem, indicating that leaf photosynthetic capacity is coordinated with stem photosynthesis on a seasonal timescale. These results suggest that both species tend to upregulate stem and leaf photosynthesis in synchrony to maximize total photosynthetic capacity during periods of intensive growth.

In contrast to total chlorophyll content, the chlorophyll a/b ratio showed pronounced seasonal variation in both species, with the lowest values occurring in summer. Chlorophyll b is an accessory pigment found in the light-harvesting complexes that extends the range of light absorption and transfers energy to photosystem-bound chlorophyll a. The lower chlorophyll a/b ratio is therefore characteristic of shade-adapted tissues (Aschan et al. 2001). The seasonal pattern in chlorophyll a/b ratio agrees well with the fact that leaf-bearing stems received only transmitted and reflected light, which was further attenuated within the radial path from the stem surface towards the photosynthetically active tissues in deeper stem layers (Pfanz et al. 2002, Natale et al. 2023a). The phellem is a protective tissue formed toward the outside of the stem and is composed of dead, suberized cells (Rosell 2019). Due to its thickness and arrangement, the phellem effectively attenuates light and reduces its penetration into deeper stem layers (Filippou et al. 2007). This corresponds with our observation that stems with thicker phellem showed a lower chlorophyll a/b ratio, indicating that the stem photosynthetic apparatus acclimated to reduced light penetration through this suberized layer (Kauppi 1991).

Stem water status, measured as stem relative water content or stem water potential, showed weak seasonal variability. We expected that stems would be more dehydrated during the peak of the growing season, when water is lost via transpiration, and the lower hydration status would be associated with higher stem photosynthesis, which could help facilitate external water uptake or mitigate drought-induced embolism (Liu et al. 2019). However, stem gross photosynthetic rates were positively related to stem water potential in our study, with less negative values being associated with higher photosynthetic rates. This result does not support our hypothesis that stem photosynthesis would be upregulated under less favorable xylem water conditions (Schmitz et al. 2012, Liu et al. 2019). However, it is important to note that stem water potentials remained well above the thresholds associated with significant embolism in both species throughout the growing season (Scholz et al. 2013, Li et al. 2015) and there was consequently little need for embolism reversal. Contrary to our expectations, the stems contained the least amount of water before the onset of the growing season, particularly in *P. avium*. It is likely that stems underwent freeze-induced embolism and tissue dehydration during the dormancy period, which could be partially reversed upon resumption of growth. Whether stem photosynthesis contributed to stem rehydration and embolism removal after winter dormancy, as reported for *Salix matsudana* (Liu et al. 2019), requires further research.

## Conclusions

Stem photosynthesis plays an important role in tree carbon balance and the maintenance of xylem hydraulic function, although net CO_2_ flux often remains negative in typical woody stems. Our results demonstrate an apparent seasonal pattern in both stem photosynthesis and respiration, reflecting seasonal shifts in growth activity. The results of our study should also be taken into account when modelling carbon fluxes in trees, as they highlight the strongly seasonal nature of the underlying processes. Stem photosynthesis was not directly related to stem chlorophyll content, suggesting that other factors, such as water status, changes in photochemical efficiency, or variation in light availability, also affect stem gas exchange. More manipulative experiments will be required to disentangle the complexity of the interplay between stem gas exchange and growth under variable environmental conditions.

## Acknowledgments

The authors thank Jana Francová and Oldřich Jůza for excellent technical assistance.

## Authors’ Contributions

R.J. designed the experimental setup, contributed to the measurements, and primarily contributed to the manuscript writing together with L.P.; E.H. carried out most of the measurements and contributed to the data analyses; L.P. and R.P. contributed to data analysis and interpretation of the results. All authors contributed to writing and revising the manuscript.

## Data and Materials Availability

The data underlying this article will be shared on reasonable request to the corresponding author.

