## Supplementary material for "Stem photosynthesis is coordinated with seasonal growth activity in two temperate tree species": Figure S1 and S2

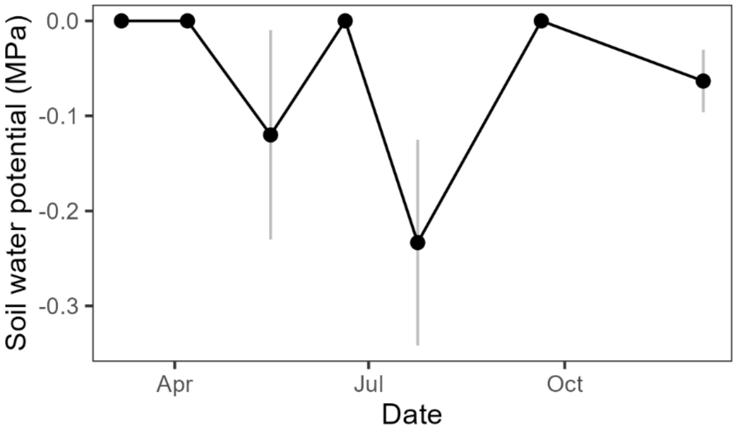


Figure S1: Seasonal course of soil water potential measured at the upper 10 cm of the soil profile within the individual sampling dates. Data are in the form means ± SE (n=3).

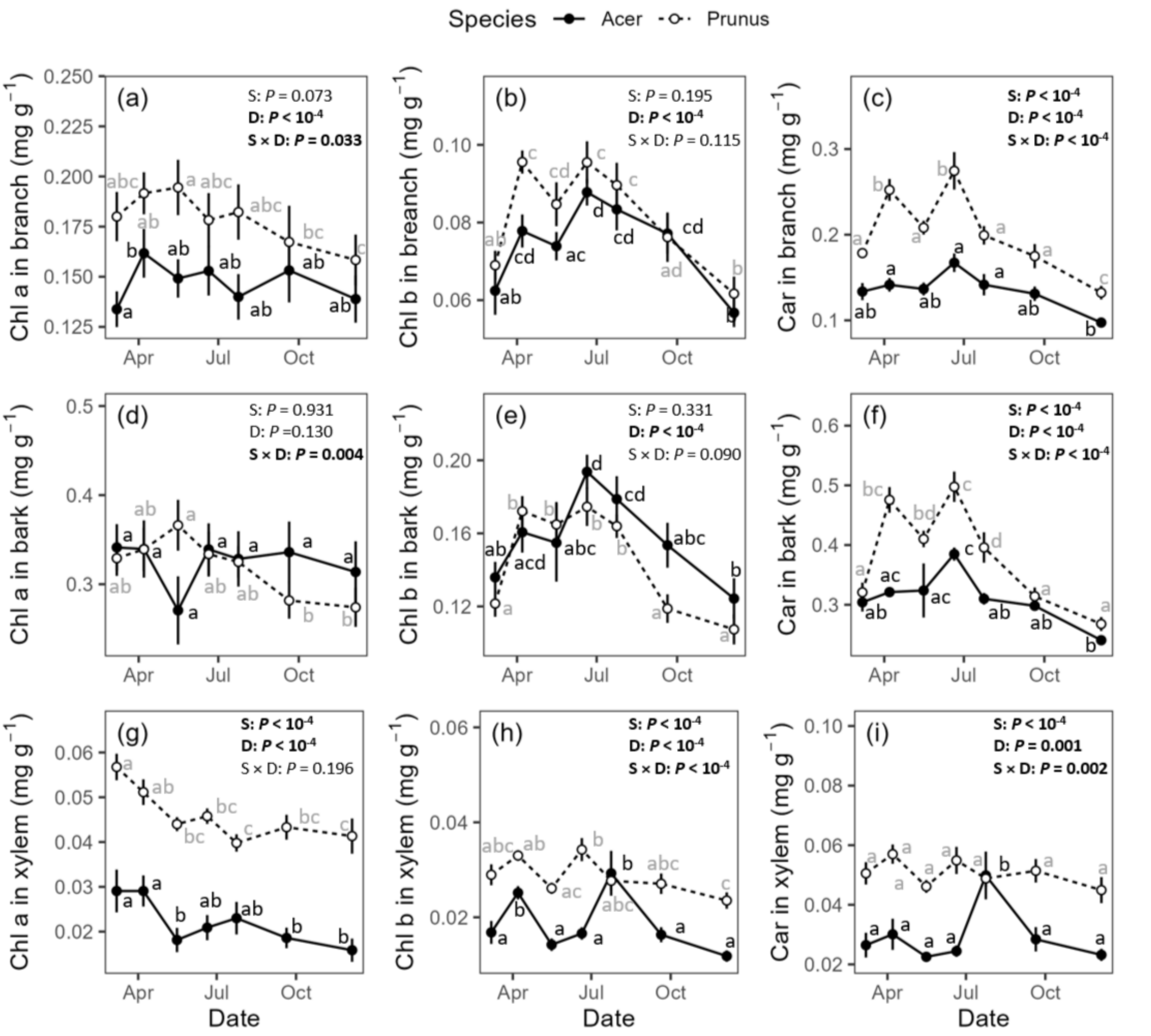


Figure S2: Seasonal variation in the chlorophyll a contents in the whole branch (a), chlorophyll b contents in the whole branch (b), carotenoid contents in the whole branch (c), chlorophyll a contents in bark (d) chlorophyll b contents in bark (e), carotenoid contents in bark (f), chlorophyll a contents in xylem (g), chlorophyll b contents in xylem (h) and carotenoid contents in xylem (i) in branches of *Acer platanoides* (closed circles) and *Prunus avium* (closed circles). A linear mixed-effects model accounting for repeated measurements on individual trees was used to evaluate the effects of species (S), sampling date (D), and their interaction (S × D). Statistically significant fixed effects are highlighted in bold. Different letters indicate significant differences within individual species among the sampling dates. Data are in the form means ± SE (n=8).
